# Revisiting equivalent optical properties for cerebrospinal fluid to improve diffusion-based modeling accuracy in the brain

**DOI:** 10.1101/2024.08.20.608859

**Authors:** Aiden Vincent Lewis, Qianqian Fang

## Abstract

**Significance:** The diffusion approximation (DA) is used in functional near-infrared spectroscopy (fNIRS) studies despite its known limitations due to the presence of cerebrospinal fluid (CSF). Many of these studies rely on a set of empirical CSF optical properties, recommended by a previous simulation study, that were not selected for the purpose of minimizing DA modeling errors.

**Aim:** We aim to directly quantify the accuracy of DA solutions in brain models by comparing those with the gold-standard solutions produced by the mesh-based Monte Carlo (MMC), based on which we derive updated recommendations.

**Approach:** For both a 5-layer head and Colin27 atlas models, we obtain DA solutions by independently sweeping the CSF absorption (*µ*_*a*_) and reduced scattering 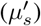 coefficients. Using an MMC solution with literature CSF optical properties as reference, we compute the errors for surface fluence, total brain sensitivity and brain energy-deposition, and identify the optimized settings where the such error is minimized.

**Results:** Our results suggest that previously recommended CSF properties can cause significant errors (8.7% to 52%) in multiple tested metrics. By simultaneously sweeping *µ*_*a*_ and 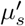,we can identify infinite numbers of solutions that can exactly match DA with MMC solutions for any single tested metric. Furthermore, it is also possible to simultaneously minimize multiple metrics at multiple source/detector separations, leading to our new recommendation of setting 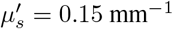 while maintaining physiological *µ*_*a*_ for CSF in DA simulations.

**Conclusion:** Our new recommendation of CSF equivalent optical properties can greatly reduce the model mismatches between DA and MMC solutions at multiple metrics without sacrificing computational speed. We also show that it is possible to eliminate such a mismatch for a single or a pair of metrics of interest.

## 1 Introduction

In the wavelength range between 600 nm and 1,100 nm, near-infrared (NIR) light can penetrate several centimeters of biological tissues with a highly scattering trajectory, as a result of relatively low tissue absorption. Multiple imaging techniques, such as functional near infrared spectroscopy (fNIRS) and diffuse optical tomography (DOT), capitalize upon this behavior to non-invasively monitor hemodynamics in cortical tissue resulting from brain activities. Similarly, in photobiomodulation (PBM) applications, clinicians use this phenomenon to deliver light to deep tissues for therapeutic purposes. Due to the highly scattering and stochastic nature of light-tissue interactions, researchers rely on quantitative modeling techniques to predict light dosages and analyze their measurements. Two widely used numerical techniques for quantitatively modeling lighttissue-interactions are the diffusion approximation (DA)^1^ and the Monte Carlo (MC) method.^2^

MC is a stochastic solver to the radiative transfer equation (RTE), a differential-integral equation known to be accurate for modeling light transport in general random media including biological tissues. MC is also relatively easy to implement and can be easily parallelized. However, the primary challenge MC faces is its high computational cost, as it requires to launch large numbers of photon packets to achieve a stable solution with acceptable stochastic noise. Over the past two decades, the widespread use of graphics processing units (GPU) has drastically reduced MC modeling time from several hours^3^ to tens of seconds.^4^ MC-based models have also been extended to accommodate increasingly complex tissue shapes, growing from infinite layered media^5^ to spatially heterogeneous voxel-based,^3, 6^ mesh-based^7^ or hybrid shape representations.^8, 9^ As a result, MC solutions have been increasingly seen in routine data analysis aside from serving its transitional role of providing gold-standard solutions.^10^

In comparison, DA solves a simplified version of the RTE by ignoring the ballistic behaviors of photons near a collimated source or in void/low-scattering regions. This results in a simpler elliptic partial different equation (PDE) that can be conveniently solved using numerical techniques such as finite-element (FE) or finite-difference (FD) methods. Typical solution time for a DA forward solution using an FE solver, such as NIRFAST^11^ or Redbird,^12^ is on the scale of a fraction of a second. This is significantly faster than MC solutions, even with GPU accelerations, and the output is deterministic (i.e. free of stochastic noise). Because of the high computational efficiency, the DA has been actively used in DOT image reconstructions especially when arrays of sources and detectors are used. However, when modeling light transport in the brain, a number of previous studies had demonstrated that the presence of low-scattering tissues such as cerebrospinal fluid (CSF) can produce erroneous solutions.^13–18^ A number of hybrid methods have been proposed to properly model voids and low-scattering tissues, such as CSF, lung, synovial fluid, cysts etc, however, these methods have received only limited adoption due to increased complexity.

The CSF layer is generally known to have low scattering, however, its literature absorption (*µ*_*a*_) and reduced scattering coefficients 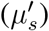 show a wide range of values, ranging between 0.0004 mm^−1^ and 0.004 mm^−1^ for *µ*_*a*_,^19, 20^ and between 0.001 mm^−1^ and 3.0 mm^−1^ for 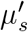 ^10, 21^ due to diverse measurement methods and modeling assumptions. The CSF layer also occupies a complex anatomical space, filling primarily the subarachnoid space bounded by the arachnoid mater at the outer surface and the pia mater at the inner surface,^22^ as well as the folding space on the cerebrum surface, known as sulci. The CSF in the sulci are predominantly transparent, filling a complex folding geometry. Sulci’s widths and depths are highly dependent on locations, ranging from 0.2 to 2.3 mm for width and 5.7 to 13 mm for depth in young adults, varying further between age groups and genders.^22^ Multiple approaches for modeling CSF in the brain exist, including treating it entirely as a translucent fluid,^20, 21^ assigning separate optical properties to the subarachnoid space and sulci,^10^ and combining it with the cortical tissue using empirically derived bulk properties.^23^

A widely cited approach for extending DA in modeling light transport in the brain is described by Custo *et al*.^20^ In this work, the CSF layer is treated entirely as a diffusive medium with a recommended empirical reduced scattering coefficient of 0.3 mm^−1^ – determined by the typical inverse line-of-sight distance of the CSF layer.^20^ Because of the simplicity, this recommmendation has been widely adopted to justify the use of DA in modeling the CSF in the brain tissues.^24, 25^ Diffusion solvers such as NIRFAST^11^ and NeuroDOT^26^ also include this recommended CSF scattering property in many built-in examples for brain related data analyses, and received wide adoption among fNIRS research.^27–29^ However, most of the works utilizing this approach took the recommended values as the optimal solution without further scrutinizing the limitations on how such a recommendation was derived.

We want to highlight that it is particularly important to understand the conditions upon which the recommended CSF optical properties in Custo *et al*.^20^ were derived. First of all, this recommendation was drawn entirely based on comparing between MC solutions at varying CSF reduced scattering coefficients, instead of directly comparing between MC and DA solutions. Secondly, the chosen value 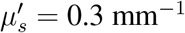 was determined as the upper-bound beyond which the MC solutions start to show large deviations from the respective ground-truth simulations; this “upper-bound” criterion was also quite different from the “optimal value” that best approximates DA with MC in the CSF that most use-cases of this recommendation commonly assumed. Thirdly, the physical quantity studied in the previous work is specifically limited to fluence and partial-path-lengths; the impact to other types of optical measurements, such as brain sensitivity – desired when solving fNIRS brain activity recovery and image reconstructions – and energy deposition – desired in many PBM related analyses – were not discussed. Lastly, the previous work only examined the impact of varying 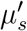; the impact of simultaneously altering CSF absorption coefficient *µ*_*a*_ was not considered.

Another potential issue in this widely cited work^20^ is the assumed “ground-truth” 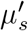 at a relatively low value of 0.001 mm^−1^. Based on the literature, this value is more closely related to *ex vivo* CSF scattering without considering subarachnoid trabeculae that reside within the CSF space. By comparing simulations with experimentally measured data, Okada *et al*.^21^ estimated that CSF in the subarachnoid space may have semidiffusive scattering values ranging from 0.16 mm^−1^ to 0.32 mm^−1^, which are significantly higher than what is used in Custo *et al*.^20^ Nevertheless, the reference CSF 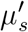 value is still under active debate. For example, Hirvi *et al*.^10^ recently suggested that the values derived in Okada *et al*.^21^ may be overestimated due to variations in CSF subarachnoid thicknesses^30^ and the fact that low-scattering regions such as sulci were not taken into account in this analysis.

Here, we would like to revisit this widely used recommendation by addressing the aforementioned limitations. Specifically, we aim to identify appropriate CSF equivalent optical properties for use in the DA to minimize the differences between DA and MC. In addition to sweeping its reduced scattering coefficient, we also allow the CSF’s absorption coefficient to change, adding a new degree-of-freedom to help reduce the model mismatch. Moreover, we expand the comparison between DA and MC to include total brain sensitivity and energy deposition, extending the new recommendation towards broader optical brain imaging/therapy techniques. Furthermore, we compare our DA and MC solutions on both a simplified 5-layered head model as well as a more complex adult brain atlas – Colin27 ^31^ – at two common wavelengths, seeking further generalization of our findings. Here we use our extensively developed mesh-based MC (MMC), known for its high accuracy among various MC solvers,^7^ with graphics processing unit (GPU) acceleration^4^ to provide the reference solution. To further remove the confounding systematic differences due to varying spatial discretization, we apply an identical set of tetrahedral meshes for use in both the FE DA solver and MMC.

In the remainder of this paper, we present our methodologies and results for minimizing DA modeling errors comparing to results obtained from MC. First, we describe the layered head and atlas anatomical models used, as well as the numerical solvers used for DA and MC respectively. Then, we describe the metrics we derive from the DA and MC solutions, such as brain sensitivity, surface fluence, and GM energy deposition; these metrics are used to quantify the errors caused by using DA in fNIRS and PBM applications. Finally, we discuss the results from both the layered and brain-atlas models, demonstrating that use of updated optical properties for CSF in DA models can significantly reduce, or even eliminate, mismatch against the “gold standard” MC models.

## 2 Methods

### 2.1 Anatomical models and tetrahedral mesh generation

We perform MC and DA simulations of light transport in two brain anatomical models frequently seen in literature: 1) a 5-layer head model and 2) Colin27 brain atlas.^32^ The 5-layered head model is created with layer thicknesses based on the average thickness of the atlas layers.^32^ Note that the scalp and skull are treated with one set of optical properties as in literature.^32^ A tetrahedral mesh of the Colin27 atlas is generated using the Brain2Mesh toolbox^32^ and is derived from the Colin27 magnetic resonance imaging (MRI) atlas, with four layers: combined scalp and skull, CSF, gray matter (GM), and white matter (WM). The physiological values for each layer, used in MC simulations serving as the ground-truth, are described in Table 1. We want to note here that all simulations in Custo *et al*.^20^ used an assumed CSF *µ*_*a*_ value of 0.004 mm^−1^. However, this value is 10× larger than the physiological *µ*_*a*_ value of 0.0004 mm^−1^ reported in Strangman *et al*.,^19^ which was also cited as the source of the optical properties. For consistency in the rest of our analysis, we use the *µ*_*a*_ values from the upstream source of Strangman *et al*.^19^ in all our reference simulations. Even though we acknowledge that the choice of the 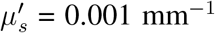 in Custo *et al*.^20^ may be problematic, as we mentioned in the Introduction section, we continue using this value in this work in order to produce meaningful comparisons with the prior work.^20^ In addition, we also repeat our analyses over an alternative 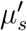 ground-truth value at 0.16 mm^−1^, as suggested by Okada *et al*.^21^This allows us to assess the robustness of our findings over significantly different assumed CSF literature values.

**Table 1.**
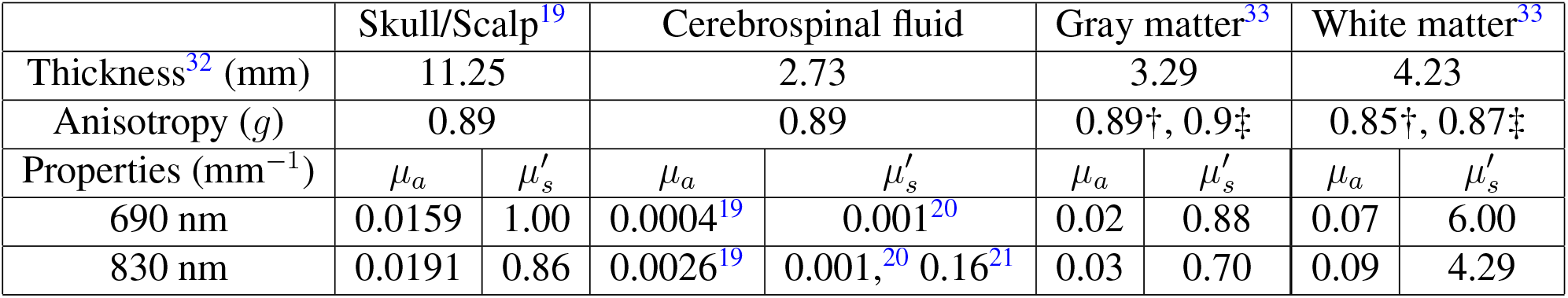
Mean tissue thicknesses, anisotropy (*g*) and assumed absorption (*µ*_*a*_) and reduced scattering coefficients 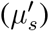, both in mm^−1^, based upon literature used to obtain the ground-truth results using Monte Carlo simulations. For anisotropy, † marks the value used for 690 nm, and ‡ for 830 nm; otherwise, the value is used for both wavelengths.

### 2.2 Forward models

We use our GPU-accelerated, mesh-based Monte Carlo (MMC) photon transport simulator^4^ – an open-source software that has been widely validated and disseminated among the biophotonics community^34–39^– to create the “ground-truth” solutions. Briefly, MMC uses tetrahedral meshes to produce MC simulations calculating fluence at every node^7^ or element. The use of tetrahedral meshes allows simulations to consider more realistic biological tissues with curved and complex boundaries. All MC solutions are produced with a relatively large number (10^9^) of photon packets to ensure stable results. For DA, we apply another in-house MATLAB toolbox named “Redbird-m” to provide diffusion solutions using a finite-element method (FEM).^12^ Redbird-m was developed from our various previous works, extending from optical breast imaging^40^ and structural-prior guided reconstructions.^41, 42^ To properly approximate a collimated source, such as a coupling fiber used in fNIRS probes, in Redbird-m, we sink sources by 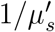 from the tissue-air boundary along the light incident direction.^43^

In all simulations reported below, both MMC and Redbird-m produce solutions over the same tetrahedral mesh generated by our Iso2Mesh mesh generator.^32^ This allows us to minimize discrepancies due to different discretization strategies. In addition, both solvers have implemented normalization methods to produce solutions that correspond to the Green’s function of the respective mathematical models, therefore, they can be directly compared at all nodal positions.

To simulate a typical fNIRS probe configuration, a pencil beam source is placed on the top surface of the chosen head model; a linear array of disk-shaped detectors with a radius of 1.5 mm, with a geodesic distance to the source ranging between 2.0 cm to 3.5 cm with an increment of 0.5 cm, are placed on one or both sides of the source. An additional near-separation detector was placed at a 0.84 cm geodesic distance from the source.

### 2.3 Metrics for accuracy assessments

To quantify the mismatch between MC and DA, we compute common metrics relevant to evaluating performance in PBM and fNIRS applications. For PBM, the total energy deposition within the GM layer (*E*_*gm*_) is computed to characterize the dosage of light energy that reaches the brain for therapeutic usage. For fNIRS, multiple metrics are used: 1) spatially resolved fluence, 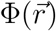, and detected fluence values, 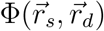, sampled at detector locations 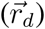 for any given source at *r*_*s*_ are extracted to assess the similarity of optical measurements between MC and DA,^13^ 2) the spatially resolved *µ*_*a*_ Jacobian, 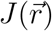,i.e. sensitivity of *µ*_*a*_ at each location 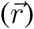,is computed to assess the loss of sensitivity caused by using DA,^44^ 3) the total brain sensitivity (*S*_*gm*_), computed by summing the Jacobian within the GM region, to estimate the overall impact of forward model accuracy to the recovery of brain hemodynamics, and finally, 4) fraction of GM sensitivity over total sensitivity (*F*_*gm*_) is used to quantify the relative impact of the forward models to fNIRS signal recovery. The definitions of each of the above metrics are detailed below.

In each of the selected head models, a forward solution is produced using MMC, by placing a pencil beam over the source position located on the skin, with an incident direction along the normal direction of the surface. In DA-based simulations, the forward solution is produced by simulating an isotropic source sunken from the skin surface by 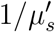 along the incident direction.^43^ Both MC and DA simulations produce forward solutions of normalized fluence, Φ, as a Green’s function defined at each spatial location 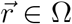 where Ω denotes the simulated domain. To calculate the total energy deposition to the brain, *E*_*gm*_, an integral of the forward fluence solution, 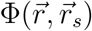, multiplied by 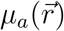 is performed within the GM region, Ω_*gm*_.

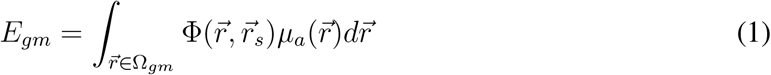

To calculate the fluence at various detectors, 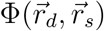, a simple linear interpolation is applied to obtain the normalized fluence at the exact coordinate 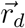 of the detector based on the forward fluence solution 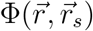 within the enclosing tetrahedron. We want to note that the fluence measurements in both MC and DA are sampled at the same spatial locations of the detectors placed along the tissue-air boundary.

We apply the adjoint method^1, 45^ to compute the Jacobian matrix in both DA and MC. This is done by multiplying the forward solution simulated from the source, 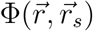, by an adjoint solution, 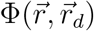, obtained by simulating a source at the location of the detector 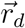. In the case of DA, we simulate both the source and the detector (as the adjoint source^46^) at their respective sunken location to mimic fiber based sources and detectors with limited numerical apertures. To compute the total brain sensitivity, *S*_*gm*_, we integrate the spatially resolved Jacobian over the entire GM region as

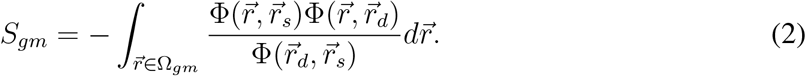

Finally, we derive the fraction of Jacobian sensitivity over the total sensitivity, *F*_*gm*_, by taking a ratio of the total brain sensitivity over the Jacobian integrated through the entire head model Ω as

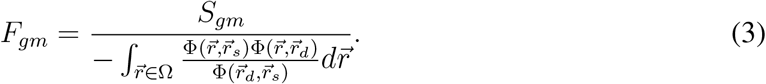

For each of the above metrics (*M*), a signed percentage error of DA relative to MC is computed as

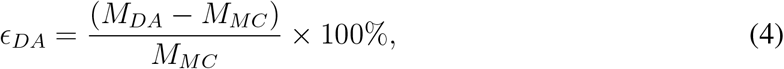

where *ϵ*_*DA*_ < 0 represents a case where DA underestimates the ground truth, and *ϵ*_*DA*_ > 0 represents overestimation.

## 3 Results

A tetrahedral mesh of 332,430 nodes and 2,015,332 elements is created using Iso2Mesh for the 5-layered head model; another tetrahedral mesh containing 151,097 nodes and 930,046 elements is produced for the Colin27 atlas. Both mesh models are shared between the MMC and Redbird-m solvers. All MMC simulations are launched on a Ubuntu 22.04.5 Linux server, running 10^9^ photon packets. This requires approximately 4.5 minutes for simulating the layered-head model and 2 minutes for the atlas model on an NVIDIA 4090 GPU. In comparison, DA solutions computed using Redbird-m are obtained using an AMD Ryzen Threadripper 3990X 64-Core processor running Linux. The DA forward simulation has a run-time around 25 seconds for the layered-head model and 15 seconds for the atlas model.

### 3.1 Assessing accuracy of diffusion approximations using literature recommendations

Contour lines in Fig. 1 compare spatially-resolved fluence (mm^−2^) computed using DA and MC across all source and detector positions; the color-maps in these plots show spatially-resolved percentage errors (*ϵ*_*DA*_), as defined in Eq. 4. In Figs. 1(a) and 1(d), we artificially set the CSF to a diffusive medium of *µ*_*a*_ = 0.0026 mm^−1^ and 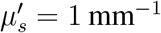 with a goal of validating Redbird-m DA solver against the reference MMC solutions. With no surprise, the Redbird-m and MMC solutions are closely aligned across the entire simulation domain in both tested head models. The excellent agreement is also indicated by the relatively uniform and low percentage errors (color shade) in most of the brain regions. In Figs. 1(b) and 1(e), we set the CSF to the anticipated physiological values 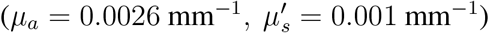.^19, 20^ The significantly elevated errors in CSF, GM and WM layers as well as at larger source-detector separations once again verify the inability of DA to model low-scattering tissues directly.

**Fig 1.**
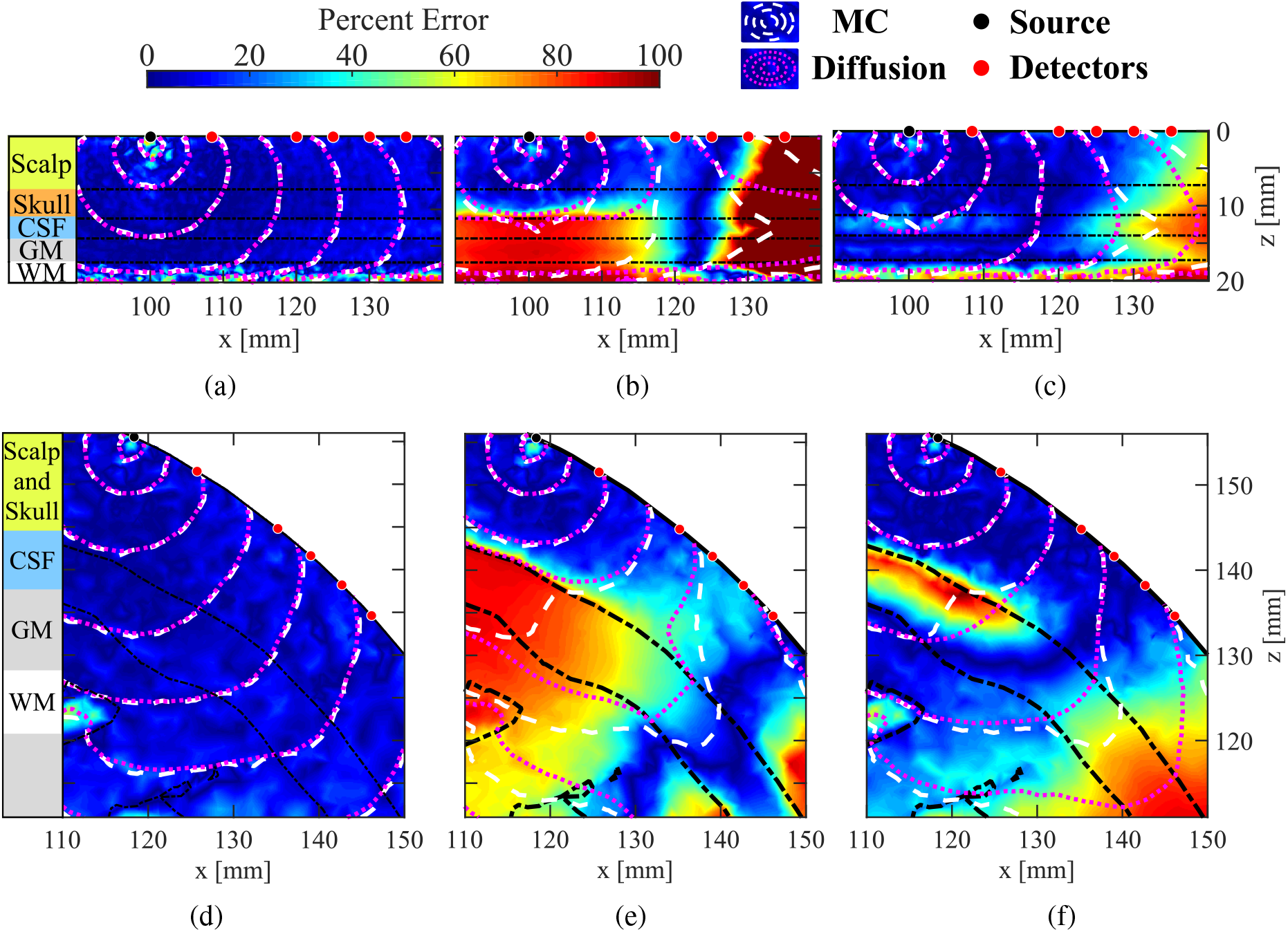
Comparison between diffusion approximation (DA) and Monte Carlo (MC) models in fluence distributions (contour lines) and percentage errors (color maps) in (a-c) a 5-layered head model and (d-f) the Colin27 atlas with a 830 nm source. The cerebrospinal fluid (CSF) optical properties are set to (a, d) a non-physiological diffusive medium of *µ*_*a*_ = 0.0026 mm^−1^ and 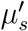 , (b, e) the assumed physiological values of *µ*_*a*_ = 0.0026 mm^−1^, 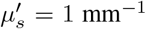 , and (c, f) the literature recommended equivalent values for DA at *µ*_*a*_ = 0.0026 mm^−1^ and 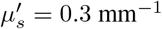 ) while MC utilizes CSF’s physiological values.

In Figs. 1(c) and 1(f), we set the CSF’s optical properties to *µ*_*a*_ = 0.0026 mm^−1^ and 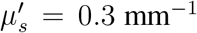 ) as recommended by Custo *et al*.^20^ It is clear from these results that this approach significantly reduces the overall mismatch between DA and MC. However, notable spatially-resolved errors ranging between 10% to 50% can still be observed within the CSF, GM and WM regions, as well as in large source-detector separations.

### 3.2 Optimization of optical properties in a 5-layer head model

To identify the optimal CSF equivalent optical properties for approximating MC simulations, in this section, we compute the DA forward solutions by sweeping CSF absorption (*µ*_*a*_) and reduced scattering coefficient 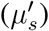 in a large search space that encompasses the typical values seen in biological tissues. The search space for *µ*_*a*_ ranges between 0 mm^−1^ and 0.04 mm^−1^ with a step size of 0.002 mm^−1^; that for 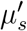 ranges between 0 mm^−1^ and 0.4 mm^−1^ in increments of 0.02 mm^−1^. The only exception is at 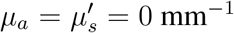 , where we had to set 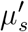 to a small value (0.001 mm^−1^) to allow the DA solver to produce valid solutions.

The signed errors (*ϵ*_*DA*_) for *E*_*gm*_, Φ, *S*_*gm*_ and *F*_*gm*_ as defined in Section 2.3, at a source-detector separation of 35 mm, are computed and plotted in Figs. 2(a) to 2(d), respectively. The color scales of all error contour plots are normalized to be between -100% and 100%, with positive errors (where DA overestimates the metric compared to MC) are shown in red and negative errors (where DA underestimates MC) are shown in blue. A red-colored square indicates the literature recommended value at *µ*_*a*_ = 0.0026 mm^−1^ and 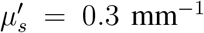 . A star is used to mark the physiological value that we used to run the MC simulation. A dashed line is plotted on each panel indicating the “zero-error contour” line where DA estimates exactly match those from MC (i.e. no error). Based on these plots, we found that the literature recommended CSF optical properties can result in −8.7% error in *E*_*gm*_, −35% error in Φ, −48% error in *S*_*gm*_ and −52% error in *F*_*gm*_ (negative errors suggest that DA underestimates MC solutions).

**Fig 2.**
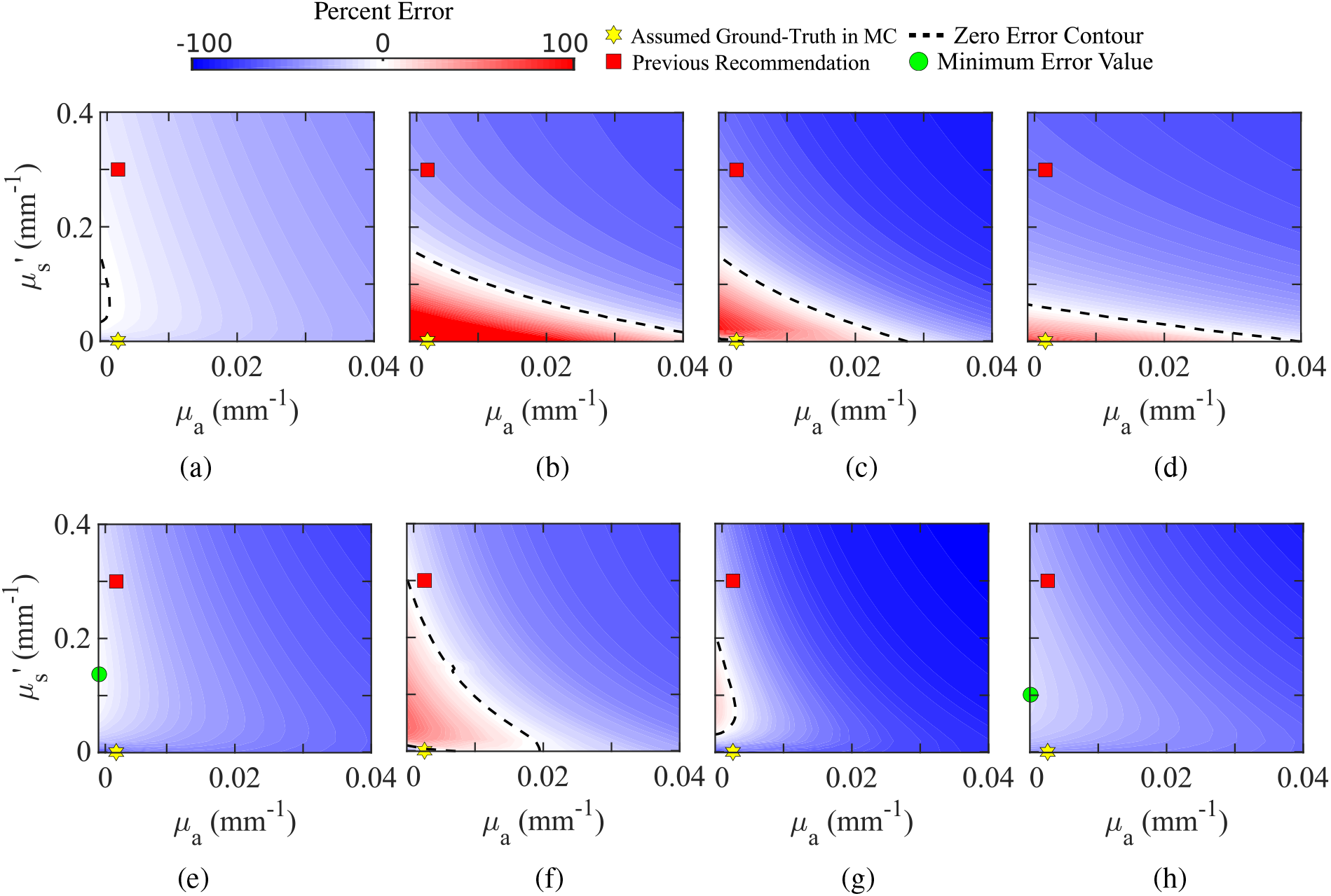
Error contour plots for DA computed at a range of CSF values compared to the MC reference solutions computed with CSF’s physiological values (*µ*_*a*_ = 0.0026 mm^−1^, 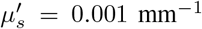 ) in a layered head model (a-d) and atlas model (e-h). We report (a, e) GM energy deposition (*E*_*gm*_), (b, f) detector fluence (Φ) at 35 mm separation, (c, g) total GM sensitivity (*S*_*gm*_) at 35 mm separation, and (d, h) fraction of GM sensitivity (*F*_*gm*_) at 35 mm separation.

We want to highlight that, for any of the given metrics, it is possible to perfectly match DA with MC values, thanks to the extra degree-of-freedom when allowing both *µ*_*a*_ and 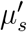 to vary simultaneously. In fact, there are an infinite number of such solutions, indicated by the continuous zero-error contour line. In other words, every combination of *µ*_*a*_ and 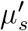 values along this line would allow the DA solution to exactly match the expected value computed by the MC model.

### 3.3 Optimization of optical properties in an adult atlas model

We repeat the above computation over the Colin27 atlas, and the error contour plots between the DA estimates (*E*_*gm*_, Φ, *S*_*gm*_ and *F*_*gm*_) at all combinations of *µ*_*a*_ and 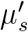 values and the respective ground-truth values obtained using MC are plotted in Figs. 2(e)-2(h). Again, we only show the plots using optical properties at 830 nm as an example; those at 690 nm look similar and are not shown.

Comparing to the plots derived from the layered head model, we found a few notable differences. First, while the error plots for Φ and *S*_*gm*_ still present the zero-error contours, *E*_*gm*_ and *F*_*gm*_ show consistent underestimation when using DA over all tested *µ*_*a*_ and 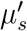 value combinations. In both cases, perfectly matching DA with MC is not possible. Instead, we indicate the combination of *µ*_*a*_ and 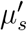 value pair where DA-to-MC error is minimized using green disk makers in Figs. 2(e) and 2(h). Another notable difference is that the contour lines of the error surfaces for the 4 selected metrics show different shapes between the layered and atlas brain models, suggesting that the choice of brain anatomical models does have notable impact to forward solutions. Nonetheless, in both cases, the overall error surfaces from all tests demonstrate a smooth monotonic trend. The desired CSF equivalent optical properties that eliminate or minimize the DA modeling errors, as either dashed line or green disks, respectively, can be readily identified from Fig. 2.

### 3.4 Simultaneously minimizing errors in two or more optical metrics

The results shown in Fig. 3 demonstrate that it is possible to completely eliminate or, in some cases, minimize modeling errors for a given metric when using DA in brain simulations by choosing a specific set of equivalent CSF optical properties. However, it is generally desirable to recommend a set of CSF equivalent optical properties that can simultaneously minimize two or more optical metrics. To achieve this goal, in Figs. 3(a)-3(b), we first show the overlay of the zero-error contour lines for *S*_*gm*_ and Φ, respectively, for the Colin27 atlas at 830 nm across 4 different separations to illustrate basic rationales when minimizing two metrics simultaneously. From Fig. 3(a), it appears that the zero-error contours for *S*_*gm*_ at various separations intersect each other in a compact region, indicated by a shaded circle. It is clear that the *µ*_*a*_ and 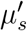 values at any intersection point of two zero-error contours is able to completely eliminate the DA modeling error for both separations. For example, *µ*_*a*_ = 0.00173 mm^−1^ and 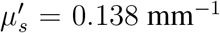 are the optimal CSF optical properties when one aims to minimize the error caused by DA for brain sensitivity at both 30 mm and 35 mm source-detector separations, with the 830 nm source. The *µ*_*a*_ and 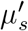 values near the center of the cluster of the intersection points, roughly located at *µ*_*a*_ = 0.0026 mm^−1^ and 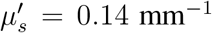 , is suitable to minimize the error for all 4 tested source-detector separations despite that it can not eliminate the error like those values at the exact intersection points. Similar optimization can be made on the error contour plots for fluence measurements Φ shown in Fig. 3(b). In this case, the intersection points are less clustered than those for *S*_*gm*_. Nonetheless, optimal CSF property settings can be found for every pair of separations.

**Fig 3.**
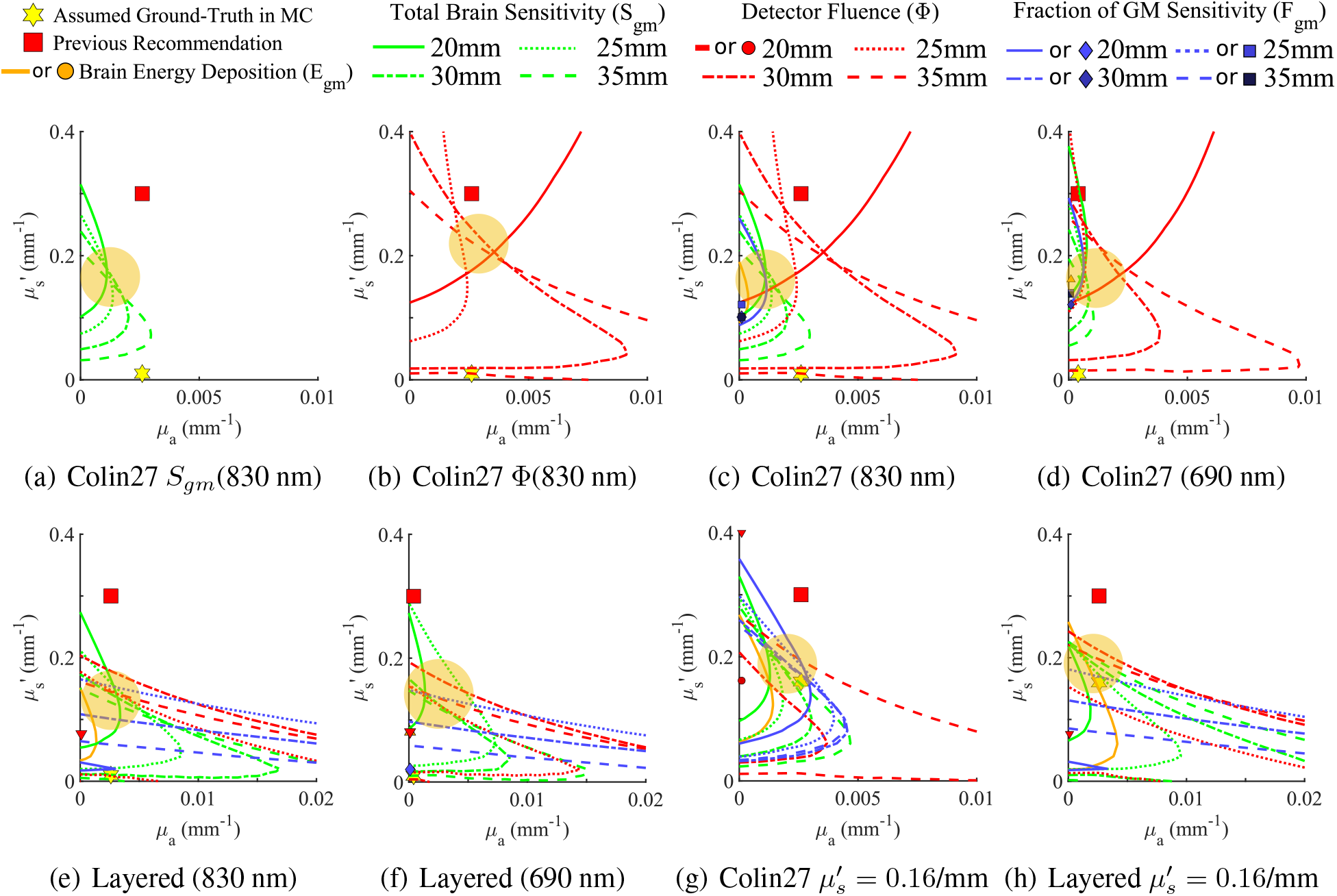
Aggregated zero-error contours and DA-to-MC error-minimizing regions between multiple metrics and head models. We show error-minimizing regions (shown as yellow-disks) for (a) GM sensitivity, (b) detector fluence, and (c) all metrics combined at 830 nm, and (d) all metrics at 690 nm in the atlas model. In the layered-head model, we show similar zero-error contour plots at (e) 830 nm and (f) 690 nm. In addition, we also include zero-error contours computed by using an alternative CSF 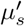 “ground-truth” value at 0.16 mm^−1^ as suggested by Okada *et al*.^21^ for (g) the atlas and (h) layered-head models at 830 nm.

In Figs. 3(c)-3(f), we show such combined zero-error contour plots for both the 5-layer (c, e) and Colin27 atlas (d, f) at either 830 nm (c, d) or 690 nm (e, f). To address the diverse CSF 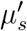 literature values, we repeat the above analysis using an alternative 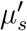 ground-truth value in the MC at 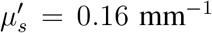 , suggested by Okada *et al*.,^21^ and report the zero-error contours for (g) the atlas and (h) 5-layer head model at 830 nm. In these plots, red lines represent errors for Φ; green lines show the errors for *S*_*gm*_, and blue lines show those for *F*_*gm*_. When a zero-error contour is not found in the search space, a marker of the corresponding color is shown marking the *µ*_*a*_ and 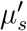 values that minimize the error of the respective metric.

It is clear that there is no single solution that can simultaneously eliminate DA modeling errors in all metrics and separations. However, we would like to highlight some general observations, from which we attempt to offer an updated recommendation for the equivalent CSF optical properties in DA.

First of all, nearly all zero-error contour lines are located below the previously recommended values (red squares), suggesting that lowering the recommended 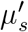 values could potentially reduce the overall modeling errors. Secondly, the brain sensitivity (*S*_*gm*_, shown in green) generally shows a more clustered intersection distribution than other metrics, with the optimal *µ*_*a*_ value in the vicinity of the physiological *µ*_*a*_ value. Thirdly, simultaneously minimizing surface fluence (Φ) at multiple separations generally requires a larger *µ*_*a*_ value, but the 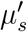 of these intersection points is comparable to those for minimizing errors of other metrics, which is about 1/2 to 1/3 of the literature recommendation. To better guide the interpretation of these findings, in Fig. 3, we draw a yellow-shaded disk on each plot indicating the rough clustered location of the zero-error contours and their intersections. When considering the alternative semi-diffusive ground-truth values for CSF suggested by Okada *et al*.,^21^ as shown in Figs. 3(g)-3(h), the error minimizing regions appear to be similar to those shown in (a)-(f), with only a slight increase in 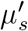.

Despite the fact that there is not a single *µ*_*a*_ and 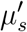 combination that could minimize all metrics, we still feel strongly that recommending a single set of *µ*_*a*_ and 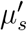 equivalent CSF properties is still quite helpful, especially considering the wide adoption of the similar recommendation from Custo *et al*.^20^ Consolidating our findings described above, we suggest to lower the 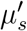 from the previously recommended 0.3 mm^−1^ to 0.15 mm^−1^ while maintaining *µ*_*a*_ to match the respective physiological values, as the error-minimizing regions tend to reside directly above the wavelength-dependent physiological *µ*_*a*_ values.

### 3.5 Verification of error reduction at optimized CSF property values

To verify that the updated recommendation of CSF equivalent properties for DA can lead to significantly lower errors over the previously recommended values, we recomputed the fluence distributions and the Jacobians for a source-detector separation of 30 mm, similar to those shown in Figs. 4(a) and 4(c), using the updated recommendations and show side-by-side comparisons in Fig. 4 before and after this optimization. The spatial distributions of the DA (magenta) and MC (white) solutions are indicated as contour lines (the closer the match, the better); the spatially-resolved percentage errors (the lower the better) of the fluence and Jacobian are also plotted as the color map in these plots.

**Fig 4.**
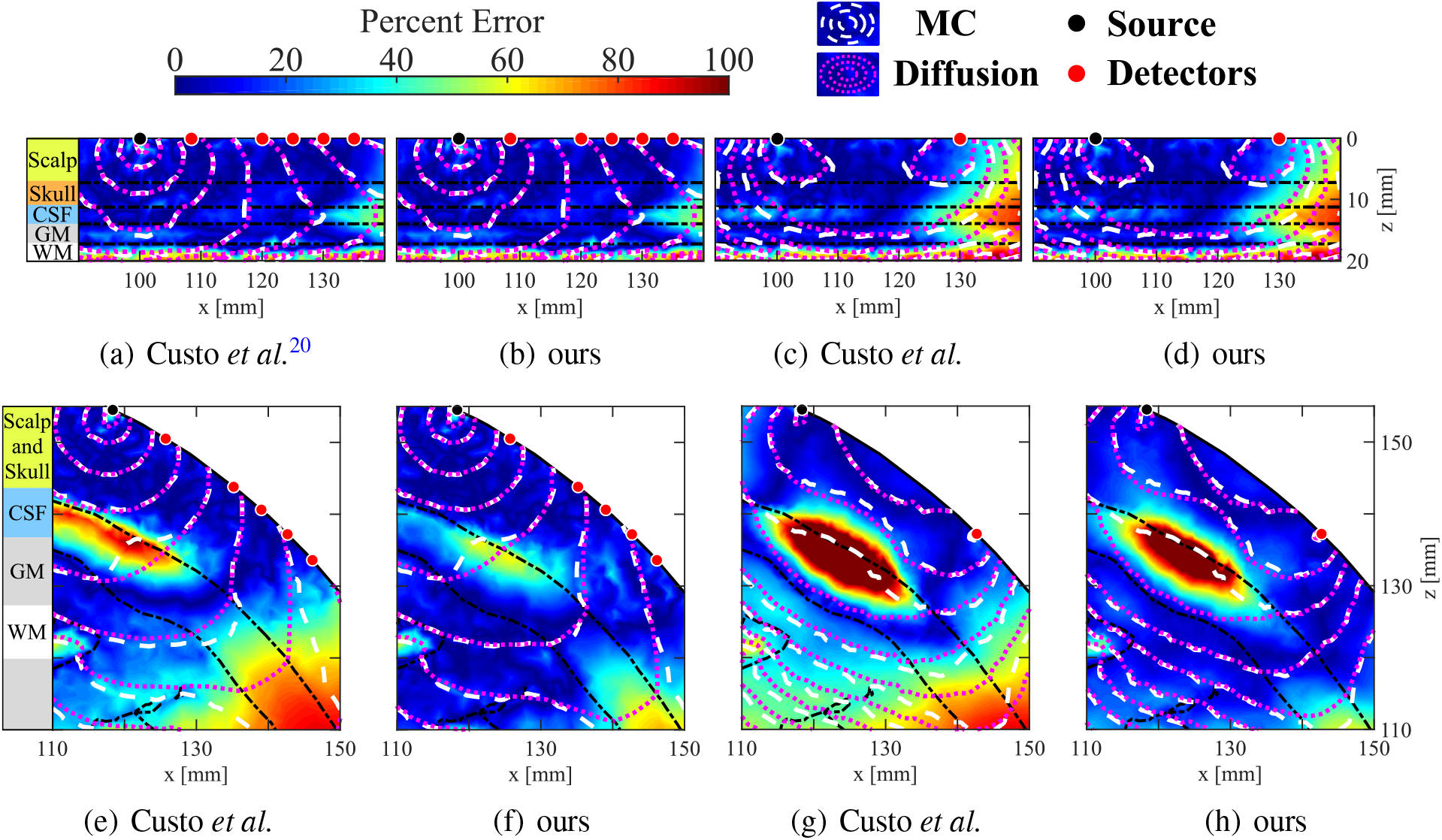
Cross sections of fluence and the 30 mm source-detector separation Jacobian in DA and MC models with a 830 nm source using previously recommended values (*µ*_*a*_ = 0.0026 mm^−1^, 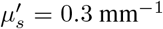 ) and our recommended values (*µ*_*a*_ = 0.0026 mm^−1^, 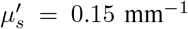 ) with absolute error (color maps) and log-scale contours of DA (magenta) and MC (white). We show a direct comparison for fluence (a, b) and sensitivity (c, d) for the layered head model, as well as fluence (e, f) and sensitivity (g, h) in the atlas model.

From fluence and Jacobian distributions in both 5-layer and atlas models, the new recommendation significantly improves the match with the ground-truth MC solutions across the domain, with particularly notable improvement in the CSF, GM and WM layers. The error distributions in all plots also become significantly more uniform across the simulated domain while shifting to-wards the low-error end. The remaining mismatch between DA and MC largely aggregates in the CSF region, while the peak error is significantly lowered when using the new recommendation. It is worth highlighting that the mismatch within the GM region has been improved dramatically. Because the GM region is particularly important in fNIRS data analysis, our updated recommendation will likely result in enhanced fNIRS analysis accuracy.

## 4 Discussions

To the best of our knowledge, this work represents the first systematic comparison between DA and MC in the handling of the low-scattering CSF tissues in brain/full-head light transport simulations. We focus on revisiting a set of equivalent CSF optical properties in DA based recommended by a widely cited work by Custo *et al*.,^20^ understanding its limitations and seeking to revise it to achieve improved modeling accuracy. Despite the relatively straightforward methodology, we believe that our updated recommendation for modeling CSF in DA is highly significant and could have a broad impact given the widespread use of the previously recommended CSF optical property values.

Based on the results presented in the above section, we want to highlight a number of key findings. First, we demonstrate that in most of the investigated optical metrics, it is possible to exactly match DA with MC when a particular optical metric is of interest. As a matter of fact, in most tested metrics, there are an infinite number of such solutions. We believe that this is a result of the additional degree-of-freedom offered by allowing both *µ*_*a*_ and 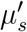 to vary. In comparison, most previous works were focused on optimizing 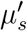 only. Secondly, a unique combination of CSF *µ*_*a*_ and 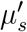 often exists to simultaneously match DA with MC when two optical metrics are considered, indicated as the intersection between any pair of zero-error contours shown in Fig. 3. Thirdly, exactly matching DA with MC solutions at more than 2 optical metrics becomes impossible, however, clusters of intersection points of zero-error curves exist and could be utilized to minimize modeling errors for a specific subset of the desired metrics. Moreover, from Figs. 2 and 3, it is clear that the previously recommended CSF optical properties tend to underestimate nearly all tested metrics when used with DA. This is indicated by the blue-colored regions where the red-square markers are located in Fig. 2 and the fact that most zero-error contours shown in Fig. 3 are below the red-square markers along the *y*-axis. Finally, using the distributions of the zero-error contours from all metrics, we reduce these complex configurations into a simple updated recommendation: we recommend lowering the CSF equivalent 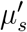 from 0.3 mm^−1^ in previous recommendation to 0.15 mm^−1^, while keeping the absorption coefficient at the physiological value.

Another advance made through this work compared to previous works is the extension from only matching the surface fluence between DA and MC to a number of fNIRS/PBM relevant metrics, including total GM sensitivity (*S*_*gm*_) and brain energy deposition (*E*_*gm*_). On the one hand, the distinctive distributions of the zero-error contour plots for each of these metrics suggest that choosing different optical metrics to optimize could lead to different optimal CSF property settings. On the other hand, comparing to the previously recommended CSF optical properties, the optimal CSF properties across various metrics all seem to require a lower 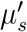 value. Similarly, our systematic benchmarks also extend the simulation domain from an atlas model in Custo *et al*.^20^ to also consider a layered brain model, which has also been frequently used in the literature.^21, 32^ While the error contour plots and zero-error curves show notable visual differences, the observations between both head models, as summarized above, are generally similar.

As we pointed out at the end of Introduction, while the assumed ground-truth CSF 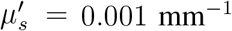 in Custo *et al*.^20^ may likely underestimate the overall CSF scattering without accounting for the presence of subarachnoid trabeculae, as suggested in Okada *et al*.,^21^ we also recognize that there is a lack of consensus in the field regarding the more appropriate alternative ground-truth 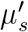 for CSF. Fortunately, from our results shown in Figs. 3(g)-3(h) using an alternative ground-truth at 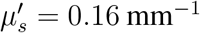 , it appears that the error-minimizing regions are largely similar to those derived using the low-scattering CSF 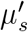 despite their 160-fold difference. This suggests that our updated recommendation can accommodate diverse assumptions regarding CSF literature values, ranging from low-scattering to sub-diffusive regimes. If one continues increasing the assumed ground-truth CSF 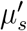 to be close to those of diffusive media, such as the upper-bound value of 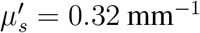 suggested by Okada *et al*.,^21^ we noticed that the error-minimizing region starts to move up in the 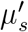 axis, and gradually approach the assumed 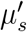 ground-truth value (not shown due to space limitations). This is anticipated because at higher 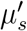 values, CSF becomes increasingly diffusive, resulting in diminished DA errors.

We recognize that there are a few limitations of this current study. First of all, our primary interest is to characterize the errors between DA with MC solutions under the influence of CSF optical properties with a continuous wave (CW) source, while assuming all other settings are identical – including the geometry of the tested brain model, the assumed optical properties of all other brain layers, even the discretization methods (i.e. meshes). One should understand that exploring variations of these other simulation assumptions, including time domain (TD) or frequency domain (FD) sources, which are beyond the scope of this work, would result in different optimal CSF optical properties. It is important to note that using DA for time domain sources may introduce additional errors.^47^ However, one could apply the same methodology as presented here to optimize CSF settings for specific application needs. Secondly, the updated CSF optical properties with 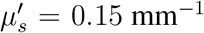 and *µ*_*a*_ at the literature physiological value is a compromise between simplicity and desired accuracy. On the one hand, the widespread use of the the previous recommendation by Custo *et al*.^20^ clearly demonstrates the need for a simple and easy-to-use approximation. On the other hand, as shown in Figs. 2 and 3, minimizing various optical metrics is a complex problem and there is not a simple solution. Thirdly, the study reported here specifically focuses on minimizing the DA-to-MC errors in selected forward and reconstruction related metrics. Despite that our updated recommendation can yield a significant reduction in error, including complete elimination in many cases. Whether such metric-wise improvement can also make a significant impact on fNIRS/PBM data analyses depends on the type of data analysis and application. In general, many fNIRS studies currently focus on relative changes between stimuli and baseline conditions. The systematic modeling errors revealed in this work may be partially alleviated due to the use of ratio-metric measurements. Regardless, for any application where the Custo *et al*.^20^ CSF DA properties were useful, the new recommendation should be readily applicable and is anticipated to achieve improved results.

## 5 Conclusion

In summary, we have systematically revisited a widely adopted recommendation on CSF equivalent optical properties to enable fast DA modeling in fNIRS data analysis with improved accuracy. By directly comparing DA with reference solutions computed by mesh-based MC, we performed comprehensive characterizations of the errors between DA and MC across various fNIRS/PBM relevant metrics, including surface fluence (Φ), total GM energy deposition (*E*_*gm*_), total GM sensitivity (*S*_*gm*_) and brain sensitivity fraction (*F*_*gm*_), at common source-detector separations across two brain models and two wavelengths. We demonstrated that by allowing simultaneous adjustments of *µ*_*a*_ and 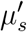, one can exactly match many single metrics between DA and MC, with an infinite number of optimal solutions along the zero-error contours. Unique optimal *µ*_*a*_ and 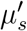 value combinations also exist among many pairs of metrics, indicated by the intersection between the two respective zero-error contours. After reviewing the overall distributions of the zero-error contours and their intersections, we found that the previously recommended CSF properties tend to underestimate many of the fNIRS/PBM relevant metrics due to their relatively high 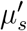 value and consequently higher attenuation.

To offer the community a convenient set of CSF optical properties for DA, we consolidated our findings across all combinations of the tested metrics, separations, brain models and wavelengths, and suggest a revised recommendation: one should set the CSF’s equivalent 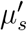 to 0.15 mm^−1^ and maintain *µ*_*a*_ at its physiological value. Compared with the previously suggested recommendation, we demonstrated that DA simulations using the revised recommendation not only significantly reduce the errors of particular fNIRS/PBM metrics, but also reduce the spatially resolved error throughout our anatomical models, especially in the GM region. In addition, we carefully described our computational protocol for this optimization; one could follow this protocol to rederive optimal CSF properties if additional constraints arise. Given the widespread use of the previously recommended CSF properties, we anticipate that our updated recommendation could generate a broad impact and improve the accuracy of fNIRS data analysis.

## Code, Data, and Materials Availability

Our open-source DA solver, Redbird-m can be accessed at https://github.com/fangq/redbird-m; our mesh-based MC solver can be accessed at https://github.com/fangq/mmc. The raw data describing relative error for all metrics and combinations of CSF *µ*_*a*_ and 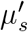 is provided in https://neurojson.org/db/cotilab/CSF_Neurophotonics_2024.

## Acknowledgments

This research is supported by National Institutes of Health (NIH) grants R01-GM114365, R01-EB026998, and U24-NS124027.

